# Small Intestinal Submucosa (SIS)-dECM Bioink with Extrusion-Induced Collagen Organization for Tympanic Membrane Tissue Engineering

**DOI:** 10.64898/2026.08.27.745367

**Authors:** Ethan Thomas John, Thirumalai Deepak, Sandhya Natesan, Lakshminath Kundanati

**Author notes:** Corresponding author: Lakshminath Kundanati, Mail ID.

## Abstract

Tympanic membrane perforations remain a common clinical problem, and although surgical intervention through tympanoplasty achieves high success rates, it is associated with donor-site morbidity, surgical complexity and limited restoration of the native radial and circumferential collagen architecture. In this study, 3D extrusion printing was utilized to create an active scaffold and attempt to promote collagen organization through shear-mediated structural alignment. An alginate-carboxymethyl cellulose (CMC) hydrogel with bovine SIS-dECM was prepared and investigated for its suitability as a bioink alternative to tympanoplasty grafts. The physiochemical, rheological and printability characteristics of the hydrogel were assessed. Successful decellularization was confirmed by histological analysis. The incorporation of the SIS-dECM into the hydrogel led to increased swelling, lower apparent viscosity, yield stress and flow stress while maintaining favourable printability and filament stability. Polarized optical microscopy was also used to study the influence of printing speed on the alignment of collagen to mimic the native tympanic membrane radial collagen architecture. Compared with the cast controls, the printed samples presented stronger birefringence signals. Biological evaluation demonstrated that the 15% dECM hydrogel exhibited the highest live cell area percentage and live/dead ratio after 48 h. In addition, the chick chorioallantoic membrane assay demonstrated that the dECM-containing hydrogels improved vascular density. The findings establish a printable, biologically active dECM bioink capable of generating bulk collagen organization through extrusion printing as a platform for tympanic membrane regeneration.

## 1. Introduction

The tympanic membrane, or eardrum, is a membranous structure separating the inner ear canal from the outer ear. It is a tri-layered structure with a medial mucosal layer, a middle fibroblast layer and a lateral epidermal layer that extends into the ear canal. The middle fibroblast layer has a unique orientation of radial and circumferentially oriented collagen fibres. These fibres play a key role in providing structural integrity and in the transmission of sound to the middle and inner ear. The tympanic membrane is an extremely thin structure, with thicknesses varying from 20 to 170 µm [1]. As such, in instances where the tympanic membrane is subjected to repeated otitis media, physical or barotrauma, it is susceptible to perforation. This occurrence is called tympanic membrane perforation (TMP). The standard practice for TMPs is surgical intervention, namely, tympanoplasty. There are several variations of tympanoplasty that are utilized on the basis of the location and extent of the perforation to the tympanic membrane. Tympanoplasties have relatively high success rates, ranging from 82% to 100% [2].

Tympanoplasties, however, are not without limitations. They require the need for autologous graft material, which the patient has to donate. Surgical expertise, persisting otorrhea, rhinitis and patient post-operative care are all contributing factors to closure of the tympanic membrane [3]. Tympanoplasties also fail at times and result in the reoccurrence of the TMP. As such, many investigations have explored the use of biomaterial-based alternatives. Owing to their high water content, tuneable mechanical properties and ability to mimic aspects of the native extracellular matrix (ECM), hydrogels have emerged as promising biomaterials for tympanic membrane repair. A variety of natural and synthetic biomaterials, including collagen [4], gelatin methacrylate (GelMA) [5,6], hyaluronic acid derivatives [7], and polycaprolactone [8], have been explored either alone or in combination for tympanic membrane tissue engineering [9–12]. Despite these advances, most currently investigated biomaterials focus primarily on achieving biocompatibility, mechanical support and favourable degradation profiles. Less attention has been directed towards reproducing the anisotropic collagen architecture characteristic of the native tympanic membrane.

Collagen organization is a defining structural feature in many biological tissue, including tendon, cartlidge, cornea, arteries and the tympanic membrane [13–16]. The orientation of collagen fibres not only governs the mechanical properties of the tissue but also influences cellular migration, proliferation and tissue remodelling. During wound healing, aligned collagen fibrils can provide contact guidance cues that direct cell migration and influence the spatial organization of the newly deposited extracellular matrix. In the tympanic membrane, radial and circumferential collagen bundles are essential for maintaining mechanical integrity and acoustic performance [17]. Disruption or incomplete restoration of this architecture has been implicated in impaired healing and chronic perforation. Therefore, biomaterials capable of recreating or directing collagen anisotropy may offer advantages for tympanic membrane regeneration.

Three-dimensional (3D) printing has been explored for the fabrication of engineered tissue constructs with precisely controlled geometries and microarchitectures. Among the available techniques, extrusion-based bioprinting has become one of the most widely explored approaches for tympanic membrane repair because of its simplicity, scalability and compatibility with a wide range of biomaterials [18]. Furthermore, an idea that has been explored is the use of the shear and extensional flows generated during extrusion to induce the alignment of fibrillar biomolecules along the direction of printing. Shear-induced alignment of collagen has been demonstrated in collagen hydrogels and composite bioinks, enabling the generation of anisotropic microenvironments that influence cell morphology and migration [19]. However, the possibility of exploiting extrusion-induced collagen alignment within decellularized extracellular matrix (dECM)-based bioinks remains relatively unexplored, particularly in the context of tympanic membrane tissue engineering.

Decellularized extracellular matrices (dECMs) are prepared with the intent of removing native cells present in tissues and organs and preserving the structure and function of the native ECM [20]. Among the numerous dECM sources, the small intestinal submucosa (SIS) has been explored owing to its composition of proteoglycans, glycosaminoglycans, and growth factors and its ease of procurement. It consequently has a pro-healing ability, desirable bioactivity and high recellularization and resorbability rates [21,22]. These factors, combined with their satisfactorily low immunogenicity [23], warrant further investigation into the use of SIS dECM for the development of bioactive bioinks capable of providing both functionally and structurally relevant collagen networks, as observed in the tympanic membrane [24].

In the present study, we developed a printable alginate-carboxymethyl cellulose (CMC) hydrogel with bovine SIS-dECM and investigated its suitability as a bioink. Owing to their favourable behaviour, mild gelation conditions and cytocompatibility, alginate-CMC hydrogels have been widely employed in extrusion bioprinting [25–27]. Here, SIS-derived dECM was incorporated to impart bioactivity and introduce collagen-rich extracellular matrix components. We hypothesized that extrusion printing could induce preferential alignment of collagenous domains within the dECM hydrogel through shear, thereby creating anisotropic architectures relevant to the native tympanic membrane. To test this hypothesis, the rheological properties, printability, collagen organization, cytocompatibility and angiogenic potential of the developed hydrogel ink were systematically investigated.

## 2. Materials and Methods

### 2.1 Materials

Sodium alginate [CAS ID: 9005-38-3], SRL, and carboxymethyl cellulose sodium salt [CAS ID: 9004-32-4], SRL, were utilized to prepare the hydrogel scaffold. The primary crosslinker was CaCl_2_ ,SRL, in its dihydrate form [CAS ID: 10035-04-8]. For the decellularization procedure reagents; Sodium hydroxide [CAS ID: 1310-73-2] SRL, sodium chloride [CAS ID: 7647-14-5] HIMEDIA, acetic acid Glacial 37.7%, peracetic acid [CAS ID: 79-21-0] HIMEDIA, for the visualization of collagen chemical reagents; Picrosirius Red [CAS ID: 2610-10-8] SRL; and formalin [CAS ID: 50-00-0] PIOCHEM was utilized.

### 2.2 Decellularization of Bovine Small Intestinal Submucosa

Bovine small intestines were procured locally from a local slaughterhouse and transported to the laboratory on ice in PBS-saline solution within three hours of sacrifice. The tissue was decellularized according to the standard optimized protocol [28,29], with an emphasis on preserving the collagenous material. Briefly, the serosa, the muscular layer and the internal mucosal layer were removed mechanically, and the small intestinal submucosa (SIS) was isolated. The SIS tissue was serially washed with 0.9% NaCl. The SIS was then subjected to two freeze thaw cycles using liquid nitrogen for 20 minutes. The tissue was then washed with 0.9% NaCl and treated with 0.1 M NaOH at room temperature for an hour, followed by another 0.9% NaCl wash. The SIS was then treated with 0.1 M ascorbic acid/0.15% peracetic acid for 36 h, followed by a 0.9% NaCl rinse again. This step was repeated with 0.15% peracetic acid in 70% ethanol for 14 h, followed by 0.15% peracetic acid in 10% H_2_O_2_ solution for 12 h. Decellularized tissues were frozen overnight at -20°C and then lyophilized at -94°C for 24 hours. The lyophilized tissue was then cryo-milled to form a dECM powder.

Decellularization efficacy was visualized through hematoxylin and eosin (H&E) staining. Decellularized and non-decellularized controls were prepared in the same way, fixed in a 10% formalin-buffered solution and embedded in paraffin blocks. Decellularized SIS samples of the tissue were sectioned with a microtome into 6 µm sections and placed on slides. These slides were immersed in xylene and ethanol solutions and then stained with H&E. Images were captured via a Labomed LX-500 light microscope.

### 2.3 Preparation of CMC-Alginate/DECM Hydrogel

To ensure the homogeneity of the hydrogel, a 2% wt/vol carboxymethyl cellulose (CMC) solution was first prepared overnight and stirred until dissolution. The next day, alginate powder was added to the CMC solution, and was mixed until it was dissolved. The 5% alginate - 2%CMC hydrogel (A5C2) was degassed via sonication with a 10-minute cycle in a 55°C water bath. The gels were then poured into 3 cc cartridges and stored at 4°C. For the preparation of the dECM hydrogels, 300 mg of dECM was dissolved in 10 ml of 0.1 M acetic acid for 48 hours at 300 rpm at room temperature. The dECM was neutralized via the addition of a few drops of 1 M NaOH. Accordingly, 10% dECM v/v and 15% dECM v/v hydrogels were prepared and subsequently referred to as A5C2d10 and A5C2d15, respectively. 100mM CaCl_2_ was utilized as an ionic crosslinker for the crosslinking of the hydrogels.

### 2.3 Physiochemical Characterization

Fourier transform infrared (FTIR) spectroscopy was carried out to identify the chemical functional groups present in the dECM hydrogel and to identify the nature of the gelation. Measurements were taken via a single reflection ATR-IR spectrophotometer (FT/IR-4X, Jasco, Japan) with an ATR (Diamond crystal) accessory. FTIR spectra were recorded in the range of 4000–500 cm^-1^ in ATR mode.

The hydrogel and decellularized SIS tissue samples were prepared for SEM by slow freezing at - 20°C. They were subsequently lyophilized at -94°C for 24 hours to prevent the formation of any artefacts during subsequent imaging. The samples were sputter coated with gold (Polaran SC7620, Quorum Technologies, UK). The microstructures of the samples were determined by scanning electron microscopy (ZEISS- ZEISS EVO 10). An accelerating voltage of 2 kV and a variable working distance between 6.88 and 12.77 mm were utilized to achieve 75x, 150x, 250x, 500x, and 1000x.

Dynamic light scattering (DLS) was utilized to characterize the particle size of the cryo-milled dECM. A HORIBA laser scattering particle size distribution analyser (LA-950V2) was utilized to determine the size of the cryo-milled dECM particles. dECM powder was dispersed in distilled water. Measurements were carried out with refractive index of water as 1.33 and that of the particle to be 1.44.

### 2.4 Swelling study

The swelling of the hydrogels was studied by immersing the crosslinked hydrogel samples in a water bath maintained at 37°C. The initial weight(W_0_) of the samples was observed, and the weights of the hydrogels were recorded at regular intervals. For the weight taken at a given time *W_t_*, the swelling ratio S can be expressed as

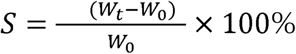

### 2.6 Rheological Characterization

The viscoelastic properties of A5C2 and A5C2-dECM were assessed with an Anton Parr Physica MCR-301 rheometer. A 25 mm diameter parallel plate setup was used, and tests were carried out with a gap of 0.52 mm. Each of the tests was carried out in triplicate. A flow sweep was carried out in the range of 1–2500 s^-1^ at a constant 19°C to understand the shear-thinning capacity of the hydrogel. The LVR (linear viscoelastic region) was identified via an amplitude sweep test, which was performed over a range of 0.01–100% strain at a fixed frequency of 1 rads^-1^ at 19°C. Within the determined LVR, a frequency sweep was conducted from 0.01–100 rad s−1 and a constant strain of 20% at 19°C to determine the storage (G’) and loss moduli (G”). The rheological data was fitted to the power law model (η=Kγ^n−1^), where *K* is the consistency index and *n* is the flow behaviour index.

### 2.7 3D Printing

An NBIL (Next big innovations lab) Trivima Pro extrusion bio-printer was used for the printing of the hydrogel scaffolds. The nozzle and cartridge temperatures were maintained at 19°C to control the self-assembly kinetics of collagen. The print bed temperature was maintained at 37°C. The constructs were printed with a printhead travel speed of 450 mm/min under pressures ranging from 1.0–1.3 Barr. Tapered needles with internal diameters of 0.41 mm (22G) and 0.26 mm (25G) were used for all print tests. The design for the prints was created manually via G-code (Marlin). To assess the printability of the hydrogel, several preliminary printing tests were carried out.

#### 2.7.1 Filament spread

To assess the diameter of the actual filament compared with the theoretical diameter of the nozzle. A simple filament spread test was conducted. A series of filaments were printed with the print speed held constant while the extrusion pressure was varied. Prior to being crosslinked, the filaments were observed under a Euromex stereomicroscope (Holland) and were analysed using the ImageFocusAlpha software. The diameters of the filaments were calculated, and the spreading factor was calculated as per the formula:

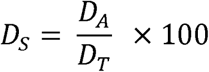

Where D*_A_* is the actual printed filament diameter and where D*_T_* is the theoretical filament diameter, which is correlated with the diameter of the nozzle.

#### 2.7.2 Filament Collapse

A filament collapse test was performed following a standard design [26,30] created with computer-assisted design. A small-filament collapse test jig was created with gaps ranging from 1 mm, 2 mm, 3 mm, 4 mm, 5 mm, and 6 mm. The design for the filament collapse jig was printed on the Bambu Labs A1 mini 3D printer using a PLA filament. Single filaments of hydrogel and dECM-hydrogel blends were printed, and their respective collapse factors were compared. A uniform air pressure of 1.4–1.5 bar was used at a feed rate of 500 mm/min, and a 25G (0.26 mm) tapered needle was used. The time taken to collapse was noted.

#### 2.7.3 Filament Fusion

A 30x30 mm test print was created, with varying spacings between filaments from 1 mm to 5 mm, thereby creating pores that increase in 1 mm intervals. Printing was carried out at 360 mm/min, 1.4 Barr and 19°C. 1-layer, 3-layer and 5-layer scaffolds were printed in triplicate. Characterizing the fidelity of the hydrogel allows a better understanding of what constructs can actually be printed with a reasonable degree of precision. The *P_r_* (printability) of the hydrogel was defined as

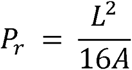

where essentially a *P_r_* value of 1 corresponds to the shape of the pore being a perfect square. values greater than 1 correspond to relative over gelation, likewise for and vice versa for values less than 1 [31]. *P_r_* was determined via ImageFocusAlpha software on the basis of images, and the printed structures were photographed via a Euromex stereomicroscope (Holland) immediately after printing without crosslinking.

### 2.8 Collagen alignment

To visualize the bulk collagen alignment present in the samples. dECM-hydrogel samples were printed with printhead travel speeds of 120 mm/min, 240 mm/min, 360 mm/min and 480 mm/min. Print controls without dECM were also printed, along with hydrogel prints that were simply cast. All the hydrogels were fixed with 2% formaldehyde for 20 minutes and then rinsed with distilled water. The hydrogels were stained with 0.1% Sirius red for one hour, and the excess stain was removed by washing with a 0.5% v/v acetic acid wash solution. The samples were consequently dehydrated in a series of increasing concentrations of ethanol. The hydrogel samples were placed on a glass slide and covered with a coverslip. Collagen was visualized with a polarized light microscope (Leica DM2500 P). The samples were viewed in transmission mode, with fixed exposure under 10x magnification. The circular stage was rotated manually to view birefringence at 0°, 45° and 90°. Images were taken and processed with Leica Application Suite (LAS) software, with only the red channel isolated to visualize Sirius red-stained collagen.

### 2.10 Live/dead Cell Assay

The cell viability and morphology were assessed by staining live cells and dead cells with calcein AM and propidium iodide (PI), respectively. The hydrogels were cast in 48-well plates, crosslinked and immersed in PBS overnight. NIH/3T3 fibroblasts were seeded onto the hydrogel scaffolds at a density of 10,000 cells/well and cultured in complete growth medium at 37°C and 5% CO_2_. After 48 hours, the samples were washed with DPBS and incubated in a staining solution containing 2 μM calcein AM and 2 μM PI. The samples were gently washed with DPBS and immediately visualized under a fluorescence microscope (Olympus IX83, Olympus).

Fluorescence images were obtained with uniform exposure and gain settings via Fiji [32]. The live-to-dead cell area ratio was calculated from the red and green channels. Image-based quantification was based on three different fields of view for each sample and from three independent replicates.

### 2.11 CAM Assay

The angiogenic potential of the hydrogels was evaluated via the chick chorioallantoic membrane (CAM) assay. Fertilized chicken eggs (embryonic day 4) were purchased from the Government Poultry Station, Potheri, Chennai, India. The eggs were incubated at 37°C and 60–65% relative humidity. On embryonic day 6, a small window was made in the eggshell under sterile conditions. The hydrogel samples were aseptically placed onto the surface of the CAM following established procedures [33]. A control group without any hydrogel was maintained in parallel. Following implantation, images were taken daily for the next 2 days (Euromex, Holland) under identical illumination and magnification settings. Angiogenesis was quantified using Fiji/ImageJ (NIH, USA). Quantitative parameters, including vessel area and vessel length, were measured within a defined region of interest surrounding the sample. Measurements were performed in triplicate for each group.

### 2.12 Statistical Analysis

All experiments were performed in triplicate unless otherwise stated, and the data are presented as the means ± standard deviations (SDs). Statistical analyses were performed via OriginPro 2025 (OriginLab Corporation, Northampton, MA, USA). Comparisons between multiple groups were carried out using one-way/two-way analysis of variance (ANOVA) followed by Tukey’s multiple comparisons test. Differences were considered statistically significant at p < 0.05.

## 3. Results

### 3.1 Evaluation of Decellularization

The efficacy of decellularization of the SIS sample was validated by H&E staining. Prior to decellularization, control SIS samples were removed and compared with decellularized SIS samples to ensure that any nuclear material (purple) was eliminated. Prior to decellularization (Figure 1a), nuclear material was observed to be distributed throughout the tissue. However, after decellularization (Figure 1b), complete removal of nuclear material was observed, and only the eosin-stained extracellular matrix tissue remained (pink), which is indicative of successful decellularization and retention of the extracellular matrix.

**Figure 1.**
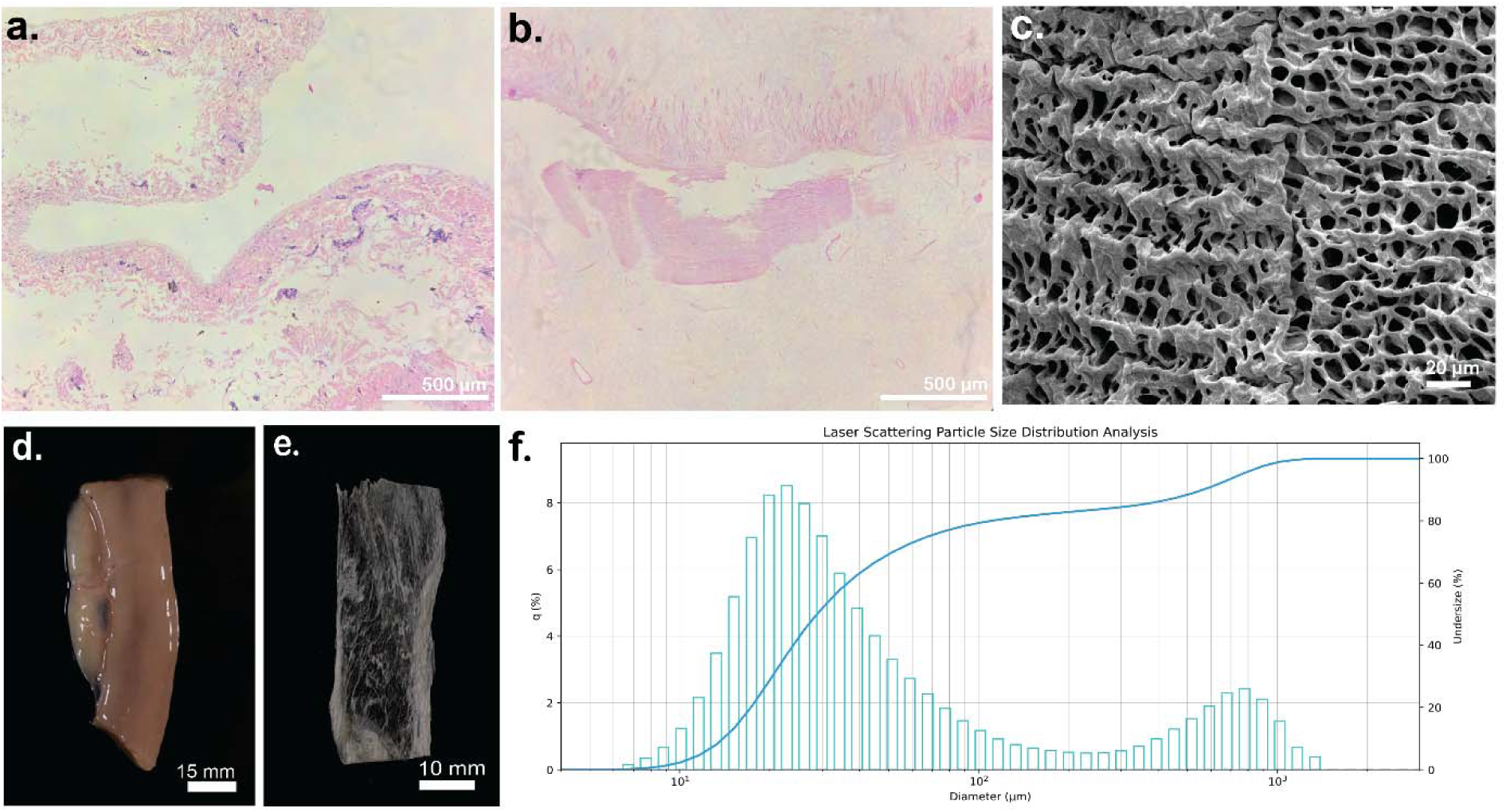
Decellularization data: a) H&E-stained section of native bovine small intestinal submucosa (nuclei in deep purple and collagenous material in pink) b) H&E-stained section of decellularized bovine small intestinal submucosa (absence of nuclei) c) SEM images of the lyophilized dECM at 1000x magnification d) image of the bovine small intestine e) decellularized and lyophilized bovine small intestinal submucosaf) SLS data indicative of the size of the dECM particles after cryo-milling

SEM imaging of the decellularized SIS revealed that the porous structure of the native extracellular matrix was largely preserved (Figure 1c). The absence of cellular debris, along with the preservation of distinct pores, is beneficial for regenerative applications, as it provides functionally relevant structural cues. Standard photographs of the SIS prior to decellularization (Figure 1d) and after decellularization (Figure 1e) are also shown. To understand the size of the milled dECM particles, static light scattering particle size analysis was conducted. This provides an estimate of the particle size prior to solubilization. The average size of the SIS particles was 28.6 ± 0.67 µm, whereas the D10 and D90 values were 13.77 µm and 472.28 µm, respectively.

### 3.2 Physiochemical Properties

#### 3.2.1 Hydrogel Swelling Study

The swelling behaviour of the hydrogels was noted over a period of 48 h (Figure 2a).All the hydrogel variants showed a marked increase in swelling between 1 and 24h. Amongst the hydrogels containing dECM, A5C2d10 and A5C2d15 slightly increased swelling. A5C2d10 exhibited swelling ratios of 81.36% and 87.6% at 24 and 48 h, respectively, whereas A5C2d15 had slightly higher swelling ratio of 96.6% and 90.36% at the same intervals. The enhanced swelling observed upon the addition of dECM suggests a resultant increase in the hydrophilicity and network architecture of the hydrogel, thereby facilitating greater water uptake.

**Figure 2.**
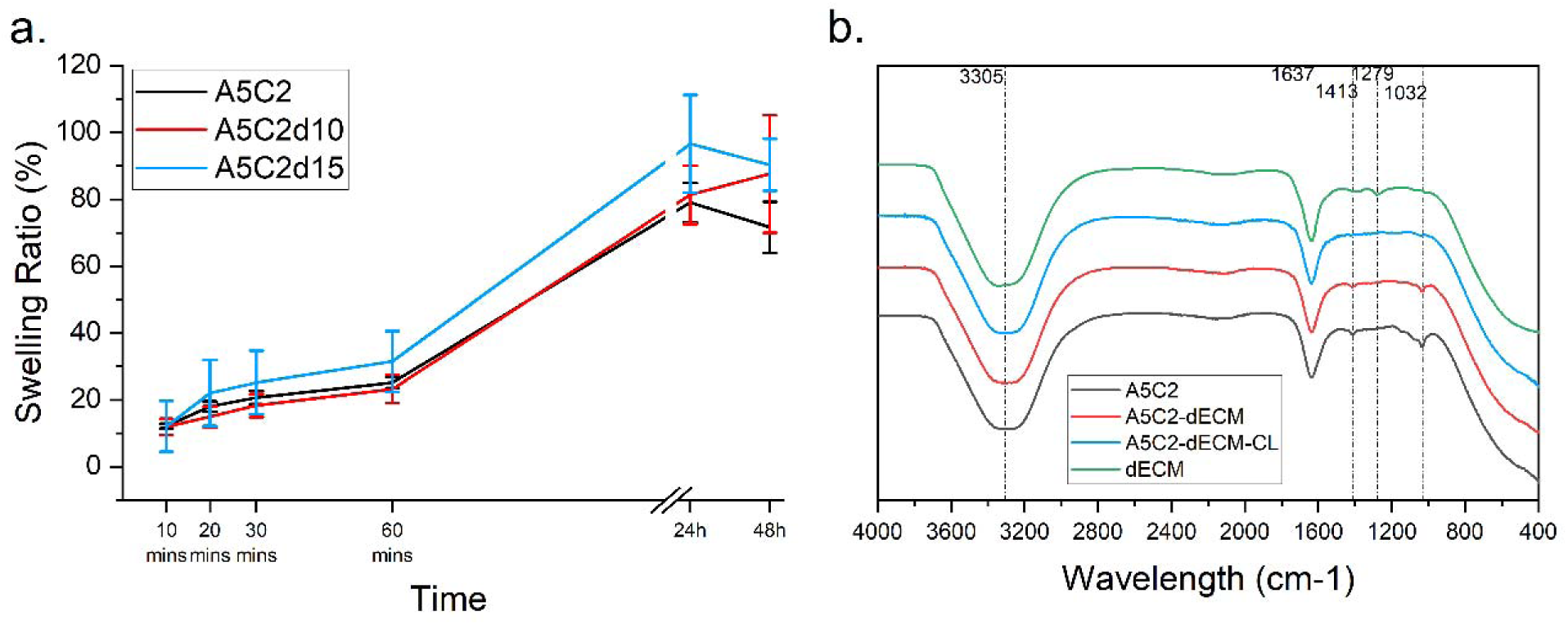
Swelling test and FT-IR. a) Swelling ratio of the hydrogels (n=3). b) FT-IR spectra of the different hydrogel compositions. dECM (in green), A5C2 (in black), A5C2-dECM (in red) and A5C2-dECM-CL (cyan), which is after crosslinking

#### 3.2.2. FTIR analysis

FT-IR analysis was carried out to understand the characteristic functional groups present in the hydrogel with and without dECM and to understand the nature of crosslinking. Distinct broad dips in transmittance are observed around the 3305 cm□¹ interval characteristic of a hydrophilic hydrogel network corresponding to -OH stretching [34]. This phenomenon was observed for all the samples owing to the aqueous nature of the hydrogels and dECM combinations. The absorption band at 1637 cm□¹ can be attributed to overlapping contributions from the amide I vibration of collagen and the bending vibration of bound water [35]. At 1413 cm□¹, COO- symmetric stretching is clearly observed in A5C2, whereas another C-O-C stretch is observed at 1032 cm ¹. Both of these peaks are greatly reduced in the crosslinked hydrogel. The latter is characteristic of polysaccharides, and this change in peak intensity can be directly attributed to the binding of Ca^2+^ to the carboxyl groups, which reduces polymer chain mobility [36]. The dECM slurry alone exhibited a band at 1279 cm ¹, corresponding to the amide III region and arising primarily from C–N stretching and N–H bending vibrations of collagen, which was absent in the alginate-CMC hydrogel alone.

### 3.3 Rheological Characteristics of the Hydrogels

Gels flowing through a dispensing needle while extruded experience shear stress. To understand how the viscosity decreases with increasing shear rate, a flow sweep was carried out on the hydrogel and hydrogel-dECM samples. The viscosity shear rate curve was fitted to a power law model, and the shear thinning nature of the polymer was noted (Figure 3a). A5C2 exhibited greater shear thinning; however, the addition of dECM to the hydrogels led to a decrease in the apparent viscosity of the dECM hydrogels. The incorporation of the dECM also significantly reduced the consistency index (*K*), from 625 for A5C2 to 232 and 217 for A5C2d10 and A5C2d15, respectively.

**Figure 3.**
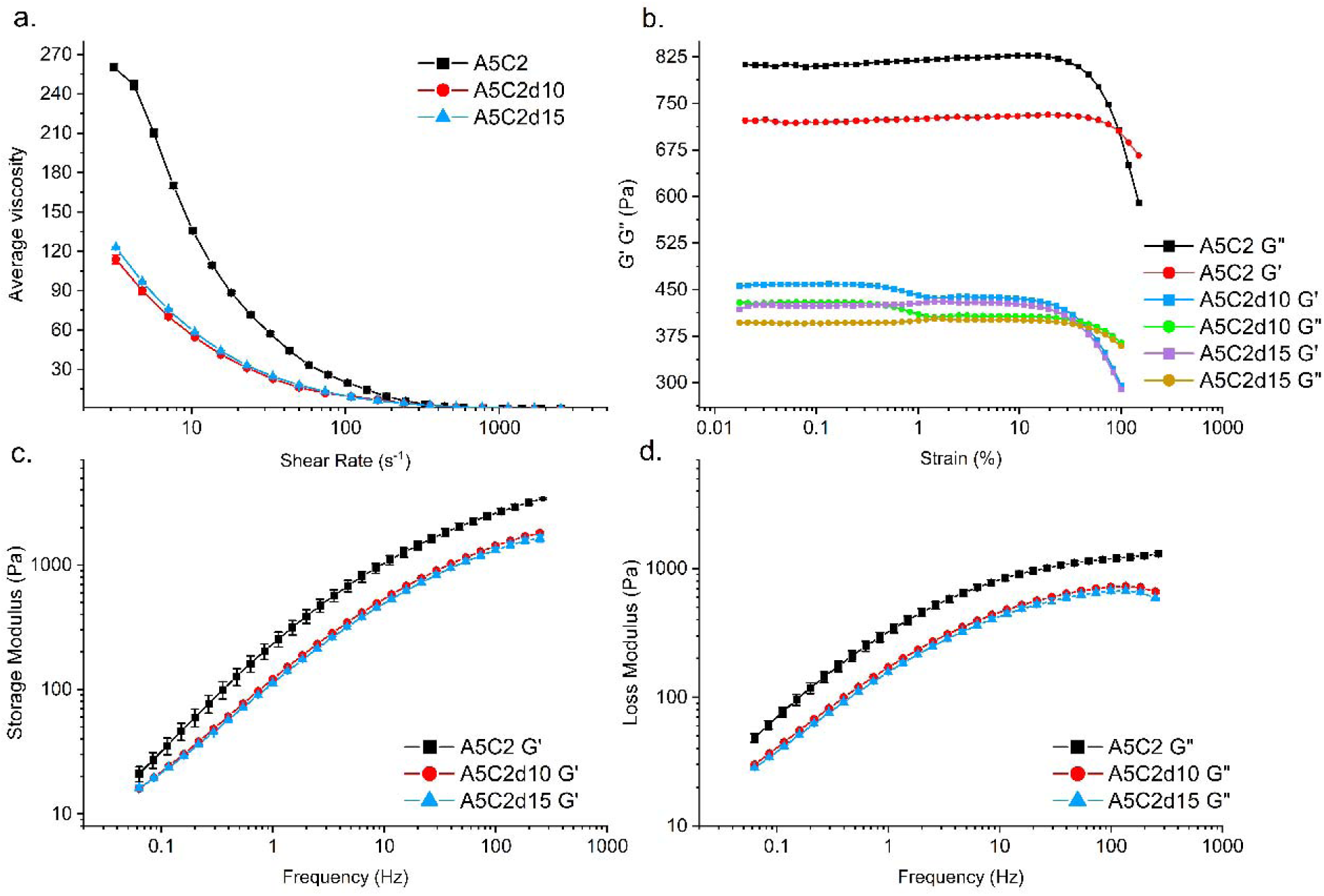
Rheological study data.a) Viscosity sweeps carried out for A5C2, A5C2d10, and A5C2 from 1 to 2500 s-1 b) amplitude sweeps carried out for A5C2, A5C2d10, and A5C2 from 0.01 to 100% strain. c) Frequency sweep storage moduli for A5C2, A5C2d10, and A5C2 from 0.01 to 100 Hz. d) Frequency sweep loss moduli for A5C2, A5C2d10, and A5C2 from 0.01 to 100 Hz.

Amplitude sweep data were obtained to identify the linear viscoelastic region (LVR) region and the storage (G’) and loss (G”) moduli (Figure 3b). The values of τ_y_ (yield stress) and τ_f_ (flow stress) were also elucidated from the amplitude sweep plot (supplementary Fig. 1). The addition of dECM to both A5C2d10 and A5C2d15 resulted in significantly lower yield stress (τ_y_) and flow stress (τ_f_) compared to A5C2. Compared across the samples, the dECM hydrogels both presented relatively lower τ_y_ and τ_f_ values. τ_y_ decreased from 33.2 Pa to 12.38 Pa for both A5C2d10 and A5C2d15. A decrease in this τ_y_ results in the requirement of lower pressure and better printability of extruded hydrogels.

The frequency sweep analysis (Figure 3c and 3d) for all the hydrogel variants revealed that, initially, the loss modulus was greater. However, it was observed that there is a crossover between G’ and G”. A5C2 exhibited this crossover at 3.53 s^-1^, whereas for the dECM variants, the crossover shifted to a higher frequency of 4.62 s^-1^. This is indicative of the fact that at frequencies lower than the crossover point, the hydrogels exhibited viscous-dominated behaviour, whereas at higher frequencies, the elastic response predominated.

**Table 1.** Rheological parameters of A5C2 and dECM hydrogels obtained from power law and amplitude sweep analyses.

| Parameter | A5C2 | A5C2d10 | A5C2d15 |
| --- | --- | --- | --- |
| $K$ | 625 | 232 | 217 |
| $n-1$ | -0.67 | -0.59 | -0.59 |
| $\tau_y$ | 33.2 Pa | 12.38 Pa | 12.38 Pa |
| $\tau_f$ | 940 Pa | 222.07 Pa | 218.5 Pa |

### 3.4 Printability of Hydrogel

The printability of the hydrogel and the dECM variants was evaluated via standard 3D extrusion-based printing tests. The incorporation of dECM altered the extrusion profile of the hydrogel and its ability to maintain structural fidelity following deposition. The filament spreading test was carried out on the A5C2 hydrogel alone and was used to optimize the printing parameters and assess the ability of the printed hydrogel filament to retain its shape. Increasing the extrusion pressure from 1 to 1.6 bar resulted in progressively greater filament spreading at all printhead travel speeds (Figure 4a), leading to increased material deposition, spreading and reduced printing fidelity. Although gels printed at 1 bar presented the lowest spreading ratios, continuous and stable filament extrusion was obtained at pressures between 1.2 and 1.4 bar while maintaining reasonable filament dimensions. Therefore, a printing pressure of 1.2 to 1.4 bar was selected as the optimal operating range, providing a balance between continuous extrusion and shape fidelity.

**Figure 4.**
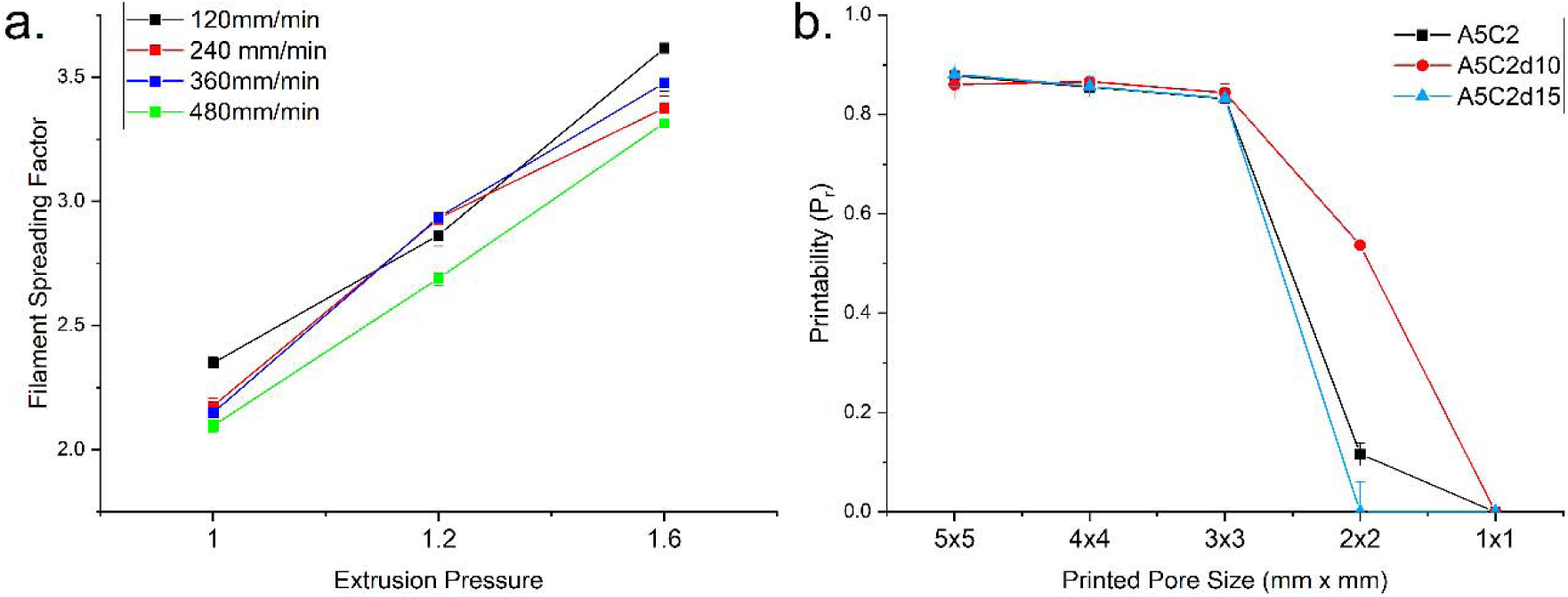
3D printing analysis. a) Filament spreading factor for different printing speeds against varying pressure. b) Printability (Pr) of the different hydrogels A5C2 (black), A5C2d10 (red), and A5C2d15 (light blue).

The filament collapse test was carried out by printing filaments across progressively increasing gap distances. All the hydrogel formulations were capable of being printed across the gaps (Figure 5a). The degree of filament sagging increased with increasing gap size for all formulations. The degree of sagging for A5C2d10 decreased, as did that for A5C2d15. The time for the filaments to collapse was noted. The A5C2-printed filament collapsed after 75 s, whereas the printed filaments containing the dECM, A5C2d10 and A5C2d15, sustained for 223 s and 214 s, respectively. The relatively favourable results can be attributed to not only exhibited lower yield and flow stresses values, but also structural contributions from the incorporated extracellular matrix.

**Figure 5.**
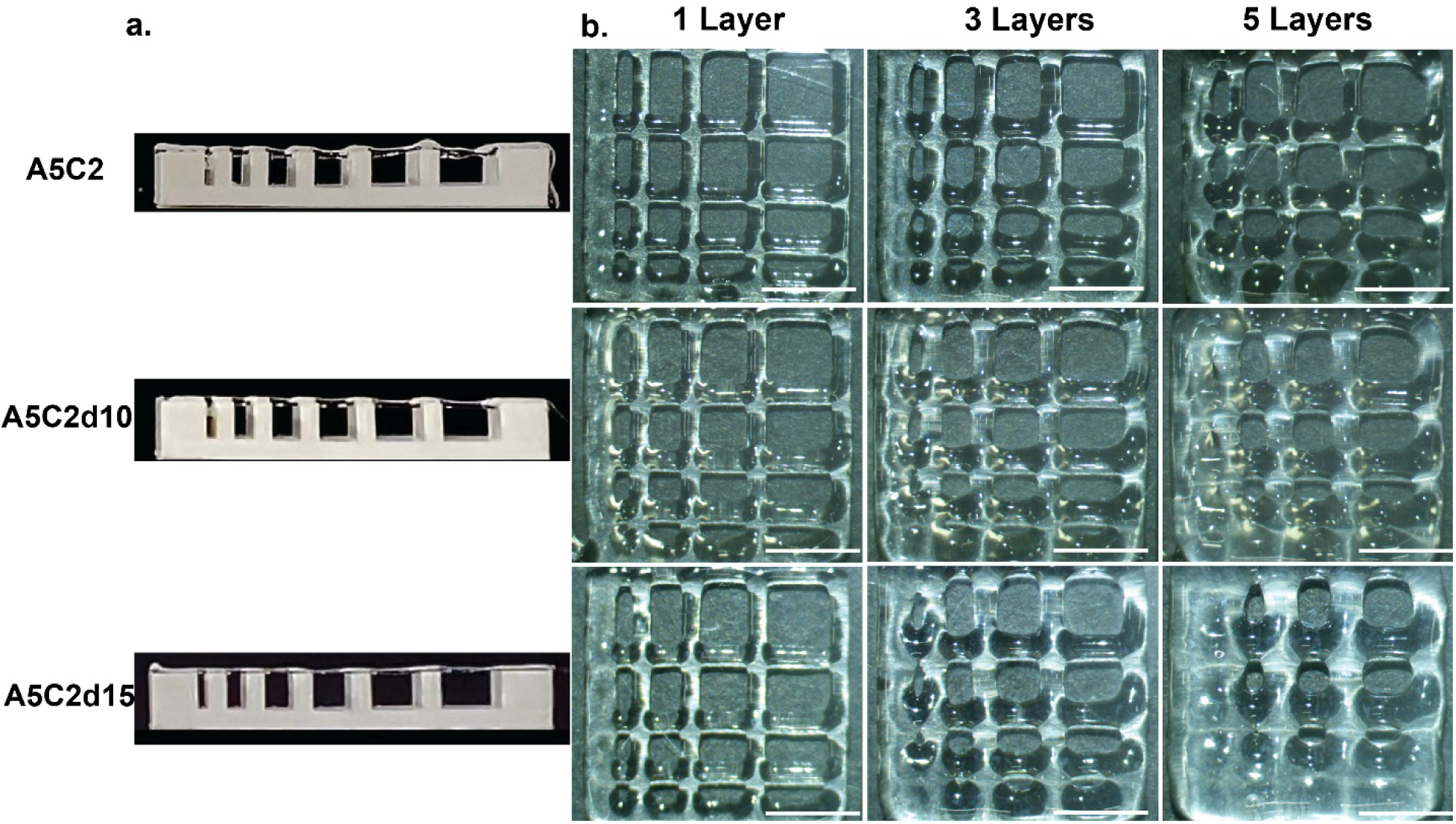
3D printing tests: a) Filament collapse test for A5C2, A5C2d10, and A5C2d15 hydrogels; b) filament fusion test with 1, 3 and 5 printed layers of hydrogel; printed hydrogel samples were taken in replicates (n=3). The scale bar at the bottom left is equivalent to 5 mm in length.

Grid structures with predefined pore geometries were printed to assess shape fidelity. A5C2 and the dECM hydrogels were capable of producing interconnected structures with pores (Figure 5b). The dECM hydrogel showed reasonable printability (P_r_) for pore sizes of 3–5 mm, with P_r_ values ranging from 0.87-0.83, which is indicative of a nearly square-like geometry, as designed for the pores. With respect to pore sizes in the range of 2 mm, a sharp drop in printability was observed overall, and none of the gels were able to consistently hold a pore size of 1 mm.

### 3.5 POM

Polarized optical microscopy was used to understand the structural organization of the collagen present in the hydrogel after extrusion printing (Figure 6). Collagen is a long rod-shaped protein with an asymmetric axis; this resulting structural anisotropy leads to observable birefringence. The birefringence pattern and intensity can be noted when picrosirius red is used. The A5C2 hydrogel exhibited small birefringent crystal domains that can be attributed to preparation artefacts. The cast dECM sample did show low-intensity birefringent sections, indicating the presence of collagen-anisotropic structures contributed by the added SIS dECM.

**Figure 6.**
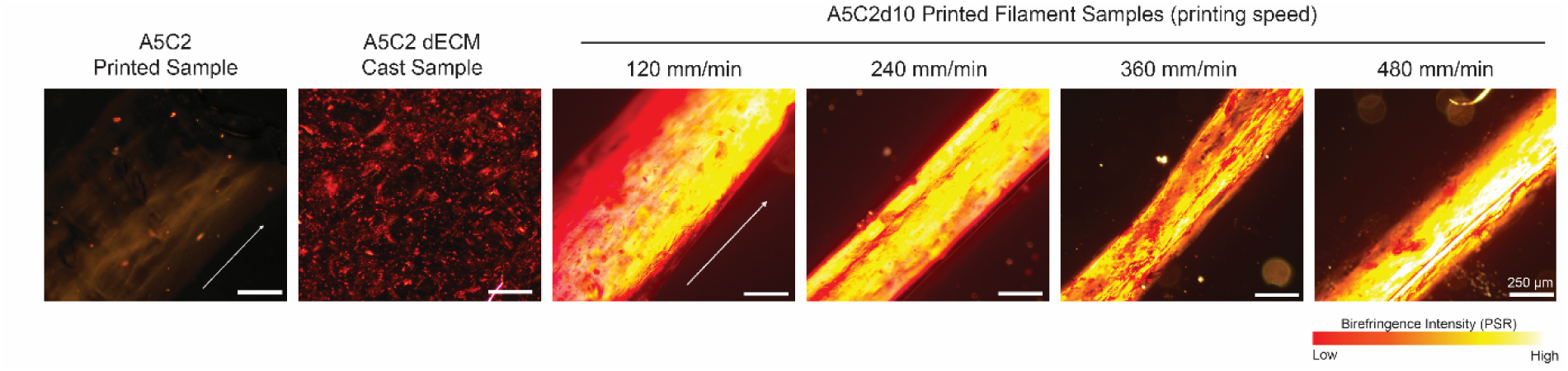
Polarized light microscopy (PLM) images of PSR-stained hydrogel formulations showing collagen birefringence and orientation. The A5C2-printed sample and A5C2-dECM-cast sample are shown as controls. Printed A5C2d10 filaments fabricated at printhead travel speeds of 120, 240, 360 and 480 mm/min exhibited progressively increased birefringence intensity and collagen alignment along the printing direction. Brighter yellow regions indicate a greater degree of collagen fibre orientation. The arrows indicate the direction of printhead travel. Scale bars = 250 µm.

The effect of the printing speed on the bulk structural organization of collagen was also studied by varying the printhead travel speed to 120, 240, 360, and 480 mm/min. All printed filament structures exhibited greater birefringence along the filament axis, with visible bright yellow regions under crossed polarizers. The anisotropic birefringence observed within the filaments suggests preferential organization of collagenous domains during extrusion that persist after crosslinking.

### 3.6 Cell Viability

The cytocompatibility of the hydrogel formulations was evaluated via live/dead staining following static seeding of L929 fibroblasts and culture for 48 h (Fig. 7a–c). Fluorescence microscopy images revealed a predominance of live cells (green fluorescence) across all formulations, with comparatively fewer dead cells (in red). Compared with A5C2C2 and A5C2d10, A5C2d15 presented a denser and more homogeneous distribution of viable cells. Quantitative image analysis further supported these observations (Fig. 7d). The live cell area increased progressively with increasing dECM content, with A5C2d15 exhibiting the highest live cell area of 32%, which was significantly greater than that of both A5C2 and A5C2d10 (p < 0.01). In contrast, the dead cell area remained low across all formulations and did not exhibit substantial differences between the groups. The live/dead ratio also increased significantly following dECM incorporation (Fig. 7e). A5C2d15 presented the highest live/dead ratio (∼12), which was significantly greater than that of both A5C2 and A5C2d10 (p < 0.01). Collectively, these results demonstrate that incorporation of dECM enhances the cytocompatibility of the hydrogel formulations and promotes fibroblast viability.

**Figure 7.**
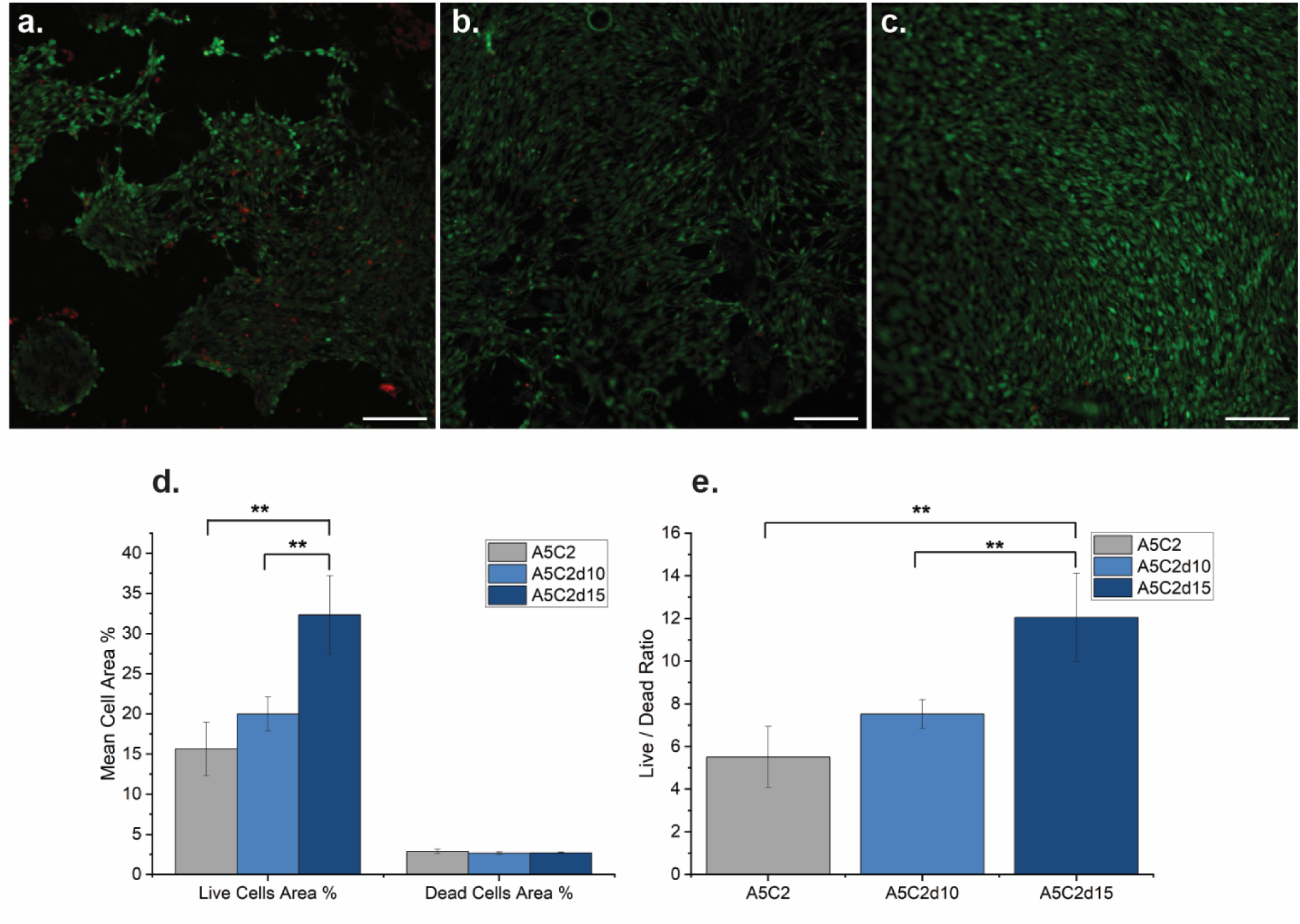
Live /dead staining of seeded NIH/3T3 fibroblasts at 48 hours. Live cells are stained green (calcein AM), and dead cells are stained red (PI). a) A5C2 hydrogel b) A5C2d10 hydrogel c) A5C2d15 hydrogel. The scale bar represents 200 µm. d) Mean cell area % of live cells and dead cells in the three samples. e) L/D ratios of cells in the three formulations. The bar plots show the means ± standard deviations generated from independent samples (n=3). Statistical analysis was performed via one-way ANOVA followed by Tukey’s post hoc test (**, p ≤ 0.01; *, p ≤ 0.05).

### 3.7 CAM Assay

The angiogenic potential of the hydrogel scaffolds was evaluated via the chick chorioallantoic membrane (CAM) assay. Representative CAM images are shown above (Figure 8). A dense network of blood vessels was present in all the embryos throughout the 2-day period. Progressive vascular remodelling was observed in all the groups.

**Figure 8.**
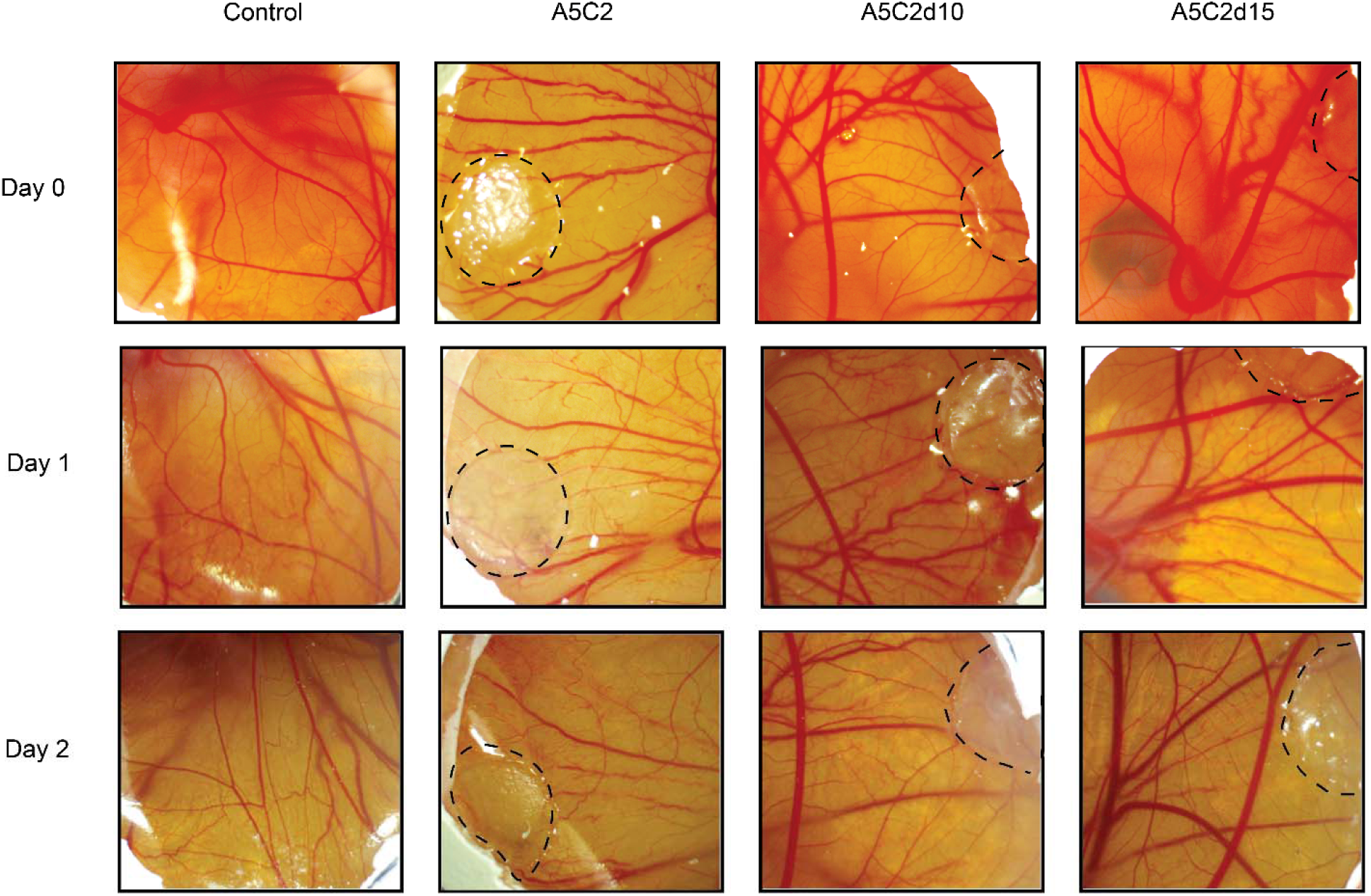
CAM assay images taken on days 0, 1, and 2 post-implantation for the control, A5C2, A5C2d10, and A5C2d15 groups. The implanted hydrogels are outlined by black dashed lines. Progressive changes in vascularization surrounding the implanted hydrogels were observed over the 2 days.

Quantitative analysis of the CAM images revealed that the vessel area fraction increased following incorporation of the A5C2 hydrogel (Fig. 9a, b). The dECM-containing hydrogels significantly increased the total blood vessel area fraction (p ≤ 0.01). However, the average diameter of the blood vessels in the control sample was greater than that in all the other hydrogel samples, with the former showing consistent values of ∼34% for the duration of the study and all the other samples showing values in the range of 14–16%.

**Figure 9.**
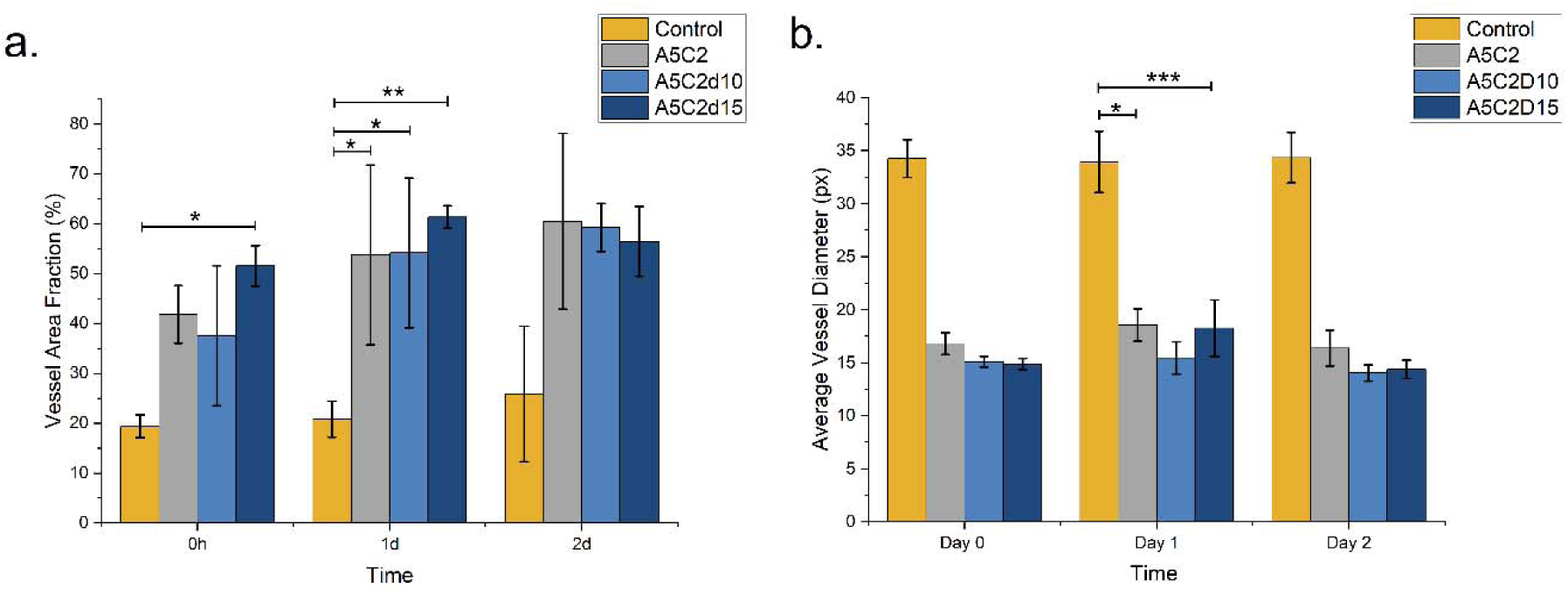
CAM results. (a) Vessel area fraction (%) of the control, A5C2, A5C2d10 and A5C2d15 groups measured at 0 h, 1 day and 2 days post-treatment. (b) Average vessel diameter (px) of the corresponding groups over the same time period. Compared with the control group, the hydrogel-treated groups presented an increased vessel area fraction accompanied by a reduction in average vessel diameter, which was indicative of the formation of a denser microvascular network. The data are presented as the means ± standard deviations from independent samples (n = 3). Statistical analysis was performed via two-way ANOVA followed by Tukey’s multiple comparisons test. Statistical significance is indicated as *p ≤ 0.05, **p ≤ 0.01, and ***p ≤ 0.001.

## 4. Discussion

### 4.1 Development of a printable SIS-dECM ink

Decellularized extracellular matrix (dECM) materials have gained considerable attention in tissue engineering owing to their ability to provide tissue-specific biochemical cues while retaining aspects of the native extracellular environment [20]. In this study, bovine small intestinal submucosa (SIS) was completely decellularized, as demonstrated by the lack of nuclear material in post decellularization imaging, while maintaining the porous architecture, as shown in the SEM images. The preservation of the collagen-rich matrix observed following decellularization is particularly important, as collagen represents a major structural component of both the SIS and the native tympanic membrane.

The swelling behaviour of the dECM-containing hydrogels indicated a slight increase in water uptake relative to that of the base hydrogel formulation. This could be attributed to an increased hydrogen bond network between the dECM and the water molecules. This increase in swelling has been previously reported with the addition of dECM [37] and RGD peptides [38]. Increased swelling, within limits, is advantageous for facilitating nutrient transport and facilitating cellular interactions within the scaffold.

### 4.2 Influence of the dECM on rheological behaviour and printability

The rheological behaviour of hydrogels is often used as a predictor of their printability, as principal parameters such as shear thinning, yield stress and damping factors directly correspond to real-life printing characteristics such as facile extrusion, flow initiation and uniformity of the printed filament, respectively [31]. Compared with A5C2 alone, the addition of SIS-dECM to the hydrogel led to a reduced apparent viscosity (v), consistency index (K), yield stress (τ_y_) and flow stress (τ_f_). The lower values of τ_y_ and τ_f_ suggest that lower extrusion forces are required to initiate and sustain flow, thereby facilitating printing under milder processing conditions.

Despite this marked reduction in viscosity and yield stress, the dECM hydrogels could exhibit improved filament stability. Conventionally, lower viscosity formulations are expected to display greater sagging and poorer shape retention. However, A5C2d10 and A5C2d15 maintained suspended filaments for substantially longer durations than the base hydrogel. A trial in which decellularized skin ECM (dsECM) was incorporated was also found to decrease filament deflection [39]. Shape fidelity testing further demonstrated that all hydrogel formulations could generate interconnected porous structures over a range of pore sizes. Importantly, the extrusion pressure was optimized for the A5C2 hydrogel, which had a relatively high τ_y_ and τ_f_ , thus the applied extrusion pressure may have resulted in greater material deposition and filament spreading for A5C2d10 and A5C2d15.

### 4.3 Extrusion-induced collagen organization and tympanic membrane repair

A defining structural feature of the native tympanic membrane is the presence of highly organized radial and circumferential collagen fibre networks [1,17]. These collagen architectures contribute significantly to the mechanical behaviour and acoustic function of the membrane [40]. Consequently, the recreation of anisotropic collagen organization represents an important objective in tympanic membrane tissue engineering.

Polarized optical microscopy was employed to investigate whether extrusion printing could influence collagen organization within the SIS-dECM hydrogel. While the cast dECM-containing samples exhibited only weak birefringence, the printed samples consistently displayed brighter birefringent regions extending along the filament axis. Furthermore, increasing the printhead travel speed resulted in a progressively stronger birefringence intensity. These observations suggest that the shear and extensional forces generated during extrusion promoted some degree of preferential organization of the collagen domains within the hydrogel [15].

It is important to note that birefringence alone does not provide direct evidence of fibril-scale collagen alignment. Rather, the present observations indicate preferential structural organization of the collagenous domains within the printed hydrogel. Nevertheless, these findings demonstrate the possibility of using extrusion printing parameters as a means of controlling microstructural organization via dECM. The use of extrusion-based printing for dECM hydrogels to generate anisotropic tissue analogues has been explored, for their use in cardiac patches, the corneal stroma and myogenic models [15,41,42] for the benefit that collagen anisotropy bears in each case. Similarly, the use of dECM-based hydrogels with controlled orientation can be explored for recreating the complex radial and circumferential collagen architectures characteristic of the native tympanic membrane.

### 4.4 Biological studies of the dECM-hydrogel

Compared with the alginate-CMC hydrogel alone, the live/dead assay demonstrated that the incorporation of SIS-dECM significantly improved fibroblast viability. The highest viability was observed in A5C2d15, which presented a significantly greater live cell area and live/dead ratio than the other formulations. These improvements likely arise from the biological complexity introduced by the extracellular matrix components. While alginate provides excellent printability and biocompatibility, it lacks many of the cell-adhesive motifs present in native extracellular matrices [43]. In contrast, the SIS dECM contains collagen and other matrix proteins capable of supporting cellular attachment and interaction, and the incorporated matrix provides a more favourable microenvironment for cell survival [22].

The CAM assay further demonstrated the biological activity of the developed bioinks. Although all the test groups exhibited vascular development over the observation period, the dECM-containing hydrogels produced a visibly denser vascular network surrounding the implantation site. Quantitative analysis confirmed increased vessel area fractions in the dECM groups relative to both the control and A5C2 formulations. The use of SIS-based dECM has been shown to result in improved vascular density in the CAM assay, which was attributed to the presence of intrinsic bFGF and vFGF [44]. While the increase in the vessel area fraction observed in the present study was modest, it followed the same overall trend reported in previous studies [45,46], suggesting that incorporation of dECM in A5C2d10 and A5C2d15 consistently promotes a proangiogenic microenvironment without markedly altering the vascular response.

Taken together, the live/dead and CAM results demonstrate that the SIS-dECM contributes to functionality beyond the structural role of the base hydrogel. The incorporation of extracellular matrix components not only improved the cytocompatibility but also enhanced the biological responses associated with tissue regeneration, highlighting the potential advantages of combining naturally derived dECM with printable hydrogel systems.

### 4.5 Implications for TM tissue engineering

Current approaches to tympanic membrane tissue engineering often focus on achieving closure of the perforation while providing temporary mechanical support during healing. However, restoring the native collagen architecture remains a significant challenge. The present study demonstrates a strategy that simultaneously addresses several important requirements for tympanic membrane regeneration, including printability, cytocompatibility, biological activity and the generation of anisotropic collagen organization.

Recent evidence suggests that tympanic membrane repair occurs through a unique epimorphic- like regenerative mechanism characterized by rapid epithelial activation, migration of progenitor populations and eventual restoration of organized collagen architecture [47]. Consequently, biomaterial strategies aimed solely at perforation closure may fail to fully recapitulate native tissue regeneration. The present findings therefore support a more structure-informed approach to scaffold design, in which extrusion-induced collagen organization is considered alongside biological performance and printability. Such strategies may ultimately contribute to the recreation of the anisotropic collagen architecture that is restored during physiological tympanic membrane regeneration.

### 4.6 Limitations and future directions

The use of dECM in 3D-printed constructs is susceptible to batch-to-batch variation, varied gelation properties, collagen content, glycosaminoglycans and growth factors [48,49]. This can consequently affect its biological performance. Future work built upon this exploratory study should investigate the reproducibility of the observed properties across batches of dECM. Another limitation is that POM provides only an indirect assessment of collagen organization and may be influenced by factors such as sample thickness, local concentration and imaging conditions. Therefore, complementary techniques capable of quantitatively resolving collagen orientation at higher spatial resolution, specifically second harmonic generation (SHG) microscopy, are needed [50]. The present study was designed as a preliminary investigation into the generation of anisotropic constructs via dECM, and future work should focus on reproducing the radial and circumferential collagen architectures characteristic of the native tympanic membrane and assessing regenerative efficacy in tympanic membrane-specific in vitro and in vivo models.

## Author Contributions

**Ethan Thomas** – conceptualization, methodology, investigation, initial draft preparation, data analysis. **Thirumalai Deepak** - methodology, investigation, data analysis**. Sandhya Natesan** performed the live/dead staining experiments and helped with related analyses. **Lakshminath Kundanati** - conceptualization, manuscript – review and editing, supervision, resources, data analysis.

## Acknowledgements

The present experimental work is funded through the institute’s NFIG grant of IIT Madras and the Ministry of Education, Govt. of India.

